# Optogenetic stimulation of the basolateral amygdala-medial entorhinal cortex pathway after spatial training has sex-specific effects on downstream activity-regulated cytoskeletal-associated protein expression

**DOI:** 10.1101/2020.04.15.042812

**Authors:** Krista L. Wahlstrom, Amanda Alvarez-Dieppa, Christa K. McIntyre, Ryan T. LaLumiere

## Abstract

Previous work from our laboratory suggests that projections from the basolateral amygdala (BLA) to the medial entorhinal cortex (mEC) are a critical pathway by which the BLA modulates the consolidation of spatial learning. Posttraining optogenetic stimulation of this pathway enhances retention of spatial memories. Evidence also indicates that intra-BLA administration of memory-enhancing drugs increases protein levels of activity-regulated cytoskeletal-associated protein (ARC) in the dorsal hippocampus (DH) and that blocking ARC in the DH impairs spatial memory consolidation. Yet, whether optical manipulations of the BLA-mEC pathway after spatial training also alter ARC in the DH is unknown. To address this question, male and female Sprague-Dawley rats received optogenetic stimulation of the BLA-mEC pathway immediately after spatial training using a Barnes maze and, 45 min later, were sacrificed for ARC analysis. Initial experiments found that spatial training increased ARC levels in the DH of rats above those observed in control rats and rats that underwent a cued-response version of the task. Optogenetic stimulation of the BLA-mEC pathway following spatial training, using parameters effective at enhancing spatial memory consolidation, enhanced ARC protein levels in the DH of male rats without affecting ARC levels in the dorsolateral striatum (DLS) or somatosensory cortex. In contrast, similar optical stimulation decreased ARC protein levels in the DLS of female rats without altering ARC in the DH or somatosensory cortex. Together, the present findings suggest a mechanism by which BLA-mEC stimulation enhances spatial memory consolidation in rats and reveals a possible sex-difference in this mechanism.

---

The basolateral complex of the amygdala (BLA) modulates the consolidation of many types of learning (Packard et al., 1994; Hatfield and McGaugh, 1999; LaLumiere et al., 2003; Miranda et al., 2003; Huff and Rudy, 2004; LaLumiere et al., 2004; McGaugh, 2004; Lalumiere and McGaugh, 2005; Roozendaal et al., 2008; Campolongo et al., 2009; Huff et al., 2013; Blank et al., 2014). The BLA maintains widespread connections with various brain regions that are selectively involved in mnemonic processes for distinct types of learning, suggesting that discrete projections are responsible for the ability of the BLA to promiscuously modulate memories (McGaugh, 2002; McIntyre et al., 2012). Consistent with this hypothesis, recent findings from this laboratory indicate that 8 Hz optogenetic stimulation of the BLA projections to the medial entorhinal cortex (mEC) selectively enhances the consolidation of spatial/context learning and impairs the consolidation of cued-response learning, a type of dorsolateral striatum (DLS)-dependent learning (Wahlstrom et al., 2018). Furthermore, optical stimulation of the BLA-mEC pathway not only increases local field potential power in the mEC but also does so downstream in the dorsal hippocampus (DH), suggesting a circuit-based mechanism by which the BLA influences hippocampal activity during the processing and consolidation of spatial information (Wahlstrom et al., 2018).

Prior work suggests that a critical mechanism for BLA influences on memory consolidation is via effects on activity-regulated cytoskeletal-associated protein (ARC) in the DH (McIntyre et al., 2005). ARC is a plasticity-associated protein that has been widely implicated in hippocampal-dependent learning and memory as a marker and effector of synaptic plasticity (Guzowski et al., 2000; McIntyre et al., 2005; McReynolds et al., 2014). *Arc* mRNA is targeted to dendrites that receive direct synaptic stimulation where it can be locally translated and interact with structural proteins to influence synaptic plasticity (Steward et al., 1998; Ploski et al., 2008). Evidence suggests that BLA activity influences hippocampal ARC protein but not mRNA expression, indicating that a post-transcriptional influence of the BLA on hippocampal ARC protein expression is a likely mechanism by which the BLA modulates memory consolidation (McIntyre et al., 2005; Ploski et al., 2008; McReynolds et al., 2014; LaLumiere et al., 2017). For example, β-adrenergic receptor activation in the BLA enhances memory consolidation for DH-dependent memories and increases DH levels of ARC but only in rats that undergo training (McIntyre et al., 2005). Similarly, post-training intra-BLA infusions of a memory-impairing dose of lidocaine significantly reduce ARC protein levels in the DH (McIntyre et al., 2005). Together, these data suggest that changes in ARC protein expression in the DH are related to observed changes in memory performance for tasks involving DH activity and that BLA activity influences ARC protein expression in the DH in a learning-dependent manner. However, whether the BLA-mEC is a circuit mechanism for influencing ARC expression in the DH is unknown.

To address this in the present study, several experiments were conducted to examine changes in ARC in the DH in relationship to learning and stimulation of the BLA-mEC pathway. Rats received optogenetic stimulation of the BLA-mEC pathway immediately following spatial training in a Barnes maze (Wahlstrom et al., 2018). Following optogenetic stimulation, rats were sacrificed for ARC protein analysis to determine the effects of posttraining BLA-mEC stimulation on ARC expression. Overall, the results suggest that such stimulation alters ARC expression in downstream regions in a complex sex- and learning-dependent manner.

## Materials and Methods

### Subjects

Male and female Sprague-Dawley rats (185-200 g and 150-175 g respectively, at time of first surgery; Envigo; *n* = 189) were used for this study. All rats were single housed in a temperature-controlled environment under a 12 h light/dark cycle (lights on at 06:00) and allowed to acclimate to the vivarium at least 3 d before surgery. Food and water were available ad libitum throughout all training and testing. All procedures used were in compliance with the National Institutes of Health guidelines for care of laboratory animals and were approved by the University of Iowa Institutional Animal Care and Use Committee.

### Surgery

Rats were anesthetized with 4% isoflurane before receiving preemptive analgesia (2 mg/kg meloxicam) and were then placed in a stereotaxic apparatus (Kopf Instruments). Surgical anesthesia was maintained with 2-3% isoflurane. All rats received virus microinjections (0.35 μL; rAAV5-CaMKIIα-hChR2(E123A)-eYFP or rAAV5-CaMKIIα-eYFP; University of North Carolina Vector Core) delivered bilaterally through a 33-gauge needle into the BLA (coordinates for males: 2.6 mm posterior and 4.9 mm lateral to bregma and 8.3 mm ventral to skull surface; coordinates for females: 2.2 mm posterior and 4.4 mm lateral to bregma and 7.4 mm ventral to skull surface). The virus injections targeted the basal nucleus of the amygdala. However, histological analysis indicated transduction of neurons throughout the basolateral complex of the amygdala, including the lateral nucleus. Thus, the present manuscript refers to the entire transduced region as the “BLA”. Four weeks later, allowing sufficient time for robust opsin expression in axon terminals, rats underwent a second surgery in which optical probes were aimed bilaterally at the mEC (coordinates for males: 7.2 mm posterior and 5.6 mm lateral to bregma and 6.5 mm ventral to skull surface; coordinates for females: 7.2 mm posterior and 4.95 mm lateral to bregma and 6.3 mm ventral to skull surface) and secured by surgical screws and dental acrylic. For each rat, a single cannula (Plastics One) that did not penetrate the skull was secured in the dental acrylic to serve as an anchor point for connection to optic leashes during optical stimulation to reduce tension on the optical fiber connections. The rats were given 1 week to recover before behavioral training.

### Optical manipulations

Optical probes were constructed by gluing an optical fiber (200 μm core, multimode, 0.37 NA) into a metal ferrule (length: 7.95 + 8.00 mm, bore: 0.230 – 0.240 mm, concentricity: <0.20 μm). The fiber extended beyond the ferrule end for implantation into tissue. The other end of the optical probe was polished and, during light delivery, connected to an optical fiber via a ceramic split sleeve. The other end of the optical fiber (FC/PC connection) was threaded through a metal leash to protect the fiber from being damaged by the rat and attached to a 1:2 splitter to permit bilateral illumination. The splitter’s single end was attached to an optical commutator (Doric Lenses) allowing free rotation of the optic leash connected to the rat. A fiber patch cable connected the commutator to the laser source (DPSS, 300 mW, 473 nm with a multimode fiber coupler for an FC/PC connection). Based on previous work, light output was adjusted to allow for 10 mW at the fiber tip (Gradinaru et al., 2009; Yizhar et al., 2011; Deisseroth, 2012; Huff et al., 2013; Wahlstrom et al., 2018), as measured by an optical power meter. Illumination was controlled by a Master-8 stimulator for all experiments involving optogenetic manipulation. The illumination was provided to rats in a separate black box holding chamber (30 cm x 30 cm x 30 cm) that contained a weighted arm attached to the outside of the chamber with the optical commutator at one end. In all cases, the comparison control was a CaMKIIα-eYFP group used for control experiments to examine the effects of illumination (and, thus, possible heating) alone. All illumination was given bilaterally.

### Behavioral training

A Barnes maze was used to investigate the consolidation of spatial and cued-response learning. The Barnes maze consisted of an exposed and elevated, brightly lit circular platform (116.8 cm in diameter) with 18 evenly spaced holes (10 cm in diameter) along the perimeter, one of which led to an escape port (see Figure 1A). The platform was covered in black vinyl in order to provide optimal contrast to the white fur of the rats for automated analysis with NOLDUS Ethovision software. Extra-maze cues consisted of specific symbols on the walls around the maze as well as the general layout of equipment in the room. NOLDUS Ethovision recording software was used to record the time to find the escape port (latency) and the time spent in each quadrant of the maze (duration).

**Figure 1.**
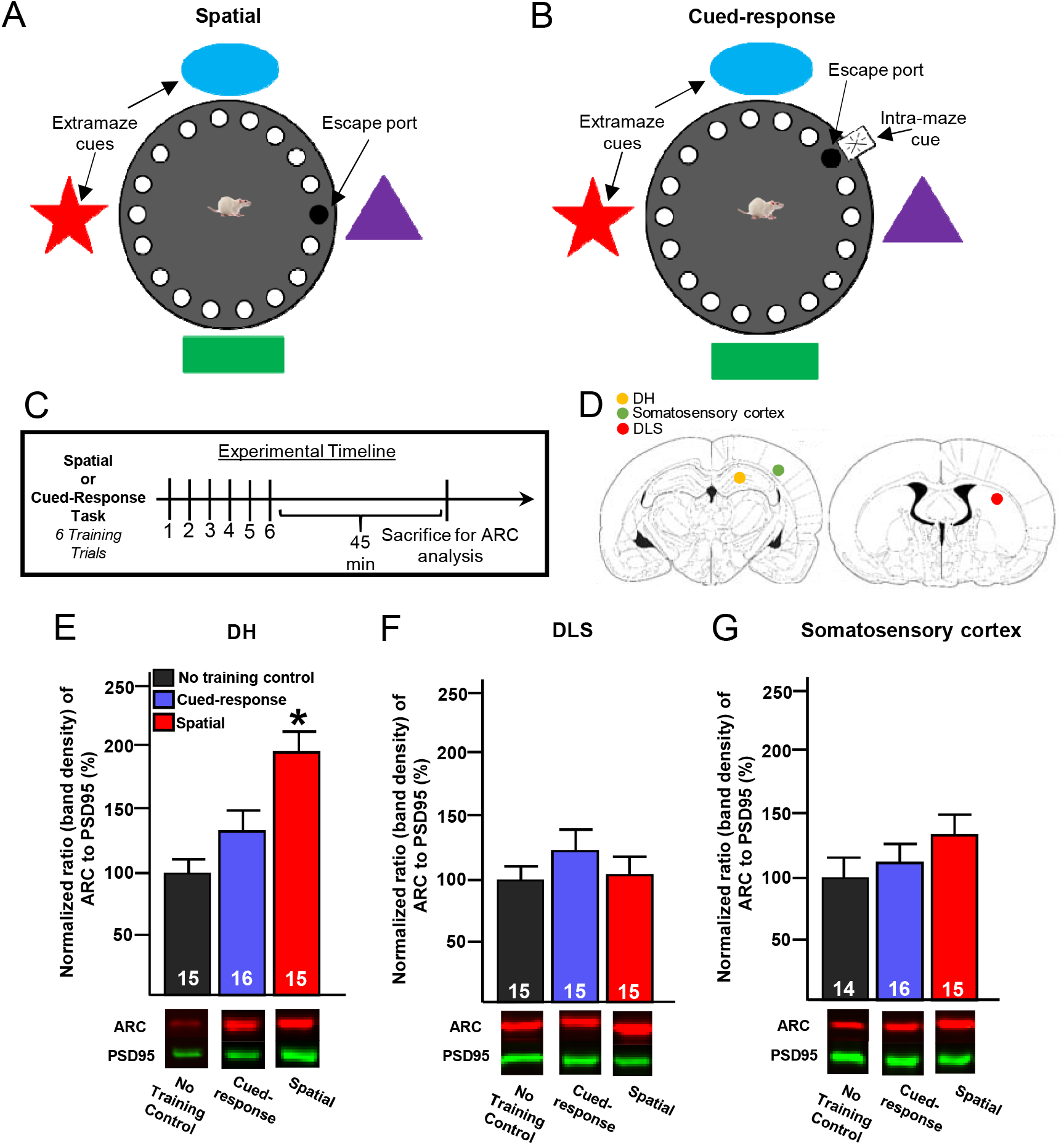
ARC protein expression effects of spatial and cued-response Barnes maze training (Experiment 1). **A and B**, Illustrations of the Barnes maze for spatial and cued-response training, respectively. For spatial training,s the escape port remained in the same location on every trial, enabling rats to use a spatial strategy to find the port. For cued-response training, a distinct intra-maze cue was attached directly to the escape port. The escape port and cue were randomly shifted to a different cardinal direction for each training trial. **C**, Experimental timeline for spatial and cued-response training. **D**, Schematic diagram illustrating the tissue punch sites in the DH, DLS, and somatosensory cortex where ARC was analyzed. **E-G**, ARC protein expression (*top*) and representative immunoblots (*bottom*) in the DH (**E**), DLS (**F**), and somatosensory cortex (**G**) for those rats that received no training, cued-response training, or spatial training. Rats that received spatial training had significantly higher ARC levels in the DH compared to no training-control rats and those that received cued-response training. There were no significant differences in ARC protein expression in the DLS or somatosensory cortex. *, p < .05 compared to no training-control and cued-response values. ARC is normalized to PSD95 by calculating the ratio band density of ARC to that of PSD95 and then expressed as a percentage of normalized control values.

For the 3 d prior to training, rats were handled individually for 1 min/d and placed in the black box holding chamber for 1 min to familiarize the rats with the environment, with the exception of the last day of handling where rats were placed in the holding chamber for 9 min. For spatial training, the escape port of the Barnes maze was maintained in the same location relative to the extra-maze cues on each trial (Figure 1A). The location was randomly chosen and counterbalanced within each group so that no particular location was associated with a single group. For cued-response training, a distinct intra-maze cue was attached directly to the escape port. The escape port and cue were randomly shifted to a different cardinal direction for each training trial (Figure 1B). Therefore, the extra-maze spatial cues could not be used to locate the escape port during cued-response training.

For both kinds of training, rats underwent multiple trials on the training day (Day 1). For each trial, the rat was placed in the center of the Barnes maze and allowed to freely explore the entire apparatus for 60 s to find the escape port and enter. If a rat entered the escape port prior to the 60 s mark, it was permitted to remain in the escape port for 30 s. If the rat did not enter the escape port within 60 s, it was placed in the escape port and permitted to remain there for 30 s. After each trial, the rat was removed from the escape port and placed in its home cage for 1 min while the maze was wiped with 20% EtOH to remove olfactory cues. This process was repeated for 6 consecutive trials (Experiment 1 spatial and cued-response strategy training) or 3 consecutive trials (Experiment 2 spatial strategy training). The number of training trials was extended for the spatial and cued-response training in Experiment 1 to ensure sufficient Barnes maze training for ARC protein expression.

In a subset of experiments (Experiment 2) retention was tested two days later (Day 3) when rats again were placed on the center of the Barnes maze and allowed to freely explore for 180 s. For this spatial version of the task, the escape port was oriented in the same direction as it had been during training on Day 1. Latency to enter the escape port and the duration spent in the target quadrant were used as the indices of retention.

### Tissue preparation

Rats were euthanized 45 min (Guzowski et al., 1999; McIntyre et al., 2005; Ramirez-Amaya et al., 2005) after the completion of spatial or cued-response Barnes maze training (Experiment 1 and Experiment 2) or 45 min after the completion of optical stimulation (Experiment 3 and Experiment 4). Animals (for ARC analysis in Experiment 1-4) were overdosed with sodium pentobarbital (200 mg/kg, i.p.), and brains were rapidly removed and flash-frozen in cold 2-methylbutane. Brains were kept at −80° C for later molecular analysis. Coronal cryosection procedures of 500 μm of thickness were taken, and the DH, DLS, and somatosensory cortex (control region) were dissected using a tissue punch kit 0.75-1.00 mm in diameter. In all three brain regions multiple tissue punches were taken from each site. The DH tissue examined was not separated into hippocampal subfields as prior works shows that the entorhinal cortex projects to all hippocampal subfields (Witter et al., 2017) and that ARC expression has similar kinetics across the entire DH (Ramirez-Amaya et al., 2005). The DLS is an area that has been previously found to be involved in cued learning in a water maze task similar to the cued-response task used herein (Packard et al., 1994) and prior work indicates that the BLA and DLS interact during memory consolidation (Goode et al., 2016). Thus, ARC expression was analyzed in the DLS for all experiments. The tissue was then stored at −80° C to be used for Western blot analysis.

### Western blots

The tissue was suspended in lysis buffer (0.1 M phosphate buffer, pH 7.4, 10% glycerol, 10% phosphatase inhibitor, and 20% protease inhibitor) and sonicated. Protein amounts were assessed using a Pierce BCA protein assay kit and spectrophotometer (Eppendorf BioPhotometer). Equal amounts of protein (15 μg) for each sample were loaded into 4-12% gradient Bis-Tris gels (Invitrogen) and separated by electrophoresis. Proteins were then transferred from the gel onto a nitrocellulose membrane by electroblotting, using an XCell SureLock wet-transfer module (ThermoFisher). The membranes were blocked with Odyssey Blocking Buffer (LI-COR Biosciences) for 1 h at room temperature. The membranes were immunoblotted with antibodies against ARC (rabbit; 1:500; Synaptic Systems) and PSD95 (rabbit; 1:1000; Cell Signaling) and incubated overnight at 4° C. Membranes were then probed with secondary antibody (IR Dye 680 LT Donkey Anti-Rabbit or IR Dye 800 CW Donkey Anti-Rabbit; LI-COR Biosciences) and incubated for 45 min at room temperature. A 1X tris-buffered saline solution containing 0.1% Tween was used to wash the membrane after incubation. Tissue from all groups was loaded into adjacent wells within a gel for fair comparison. Detection of immunoreactivity to ARC and PSD95 in each sample was determined by imaging with the Odyssey imaging system (LI-COR Biosciences) and average density of each ARC band was normalized to the PSD95 band for each sample (Alvarez-Dieppa et al., 2016) and then expressed as a percentage of normalized control values.

### Experimental design

#### Experiment 1 – ARC expression following spatial or cued-response training

Experiment 1 examined ARC protein expression in the DH, DLS, and somatosensory cortex following spatial or cued-response training. For Experiment 1, rats received either spatial or cued-response (Figure 1A and 1B) training, or were part of a no training-control group in which they were removed from their home cage and placed in the black box holding chamber (same as the illumination black box holding chamber used for optogenetic manipulations in Experiment 2-4) for 15 min. Six training trials were used in Experiment 1, as it was expected that more training trials would produce sufficient learning to identify changes in ARC expression. Following training, rats were euthanized for ARC protein analysis.

#### Experiment 2 – Post-spatial training BLA-mEC stimulation effects on ARC expression

Experiment 2 examined whether stimulating the BLA-mEC pathway after spatial training in a Barnes maze task alters ARC protein expression in downstream regions. For this experiment, rats received illumination of either ChR2 or eYFP control-transduced BLA fibers in the mEC immediately following the final spatial training trial (Figure 2A and 2B), using the following illumination parameters as previously used: 15 min of 2 s-trains of 8 Hz light pulses (pulse duration = 5 ms), given every 10 s (Wahlstrom et al., 2018). Three training trials were used in Experiment 2 to prevent ceiling effects in order to observe any enhancement in learning. Half of the animals were sacrificed following post-training stimulation to assess the effects of 8 Hz stimulation on ARC protein expression in the DH, DLS, and somatosensory cortex. The other half of the animals were given a single retention test 2 d after spatial training to verify our previously published work demonstrating that post-training stimulation enhances spatial memory.

**Figure 2.**
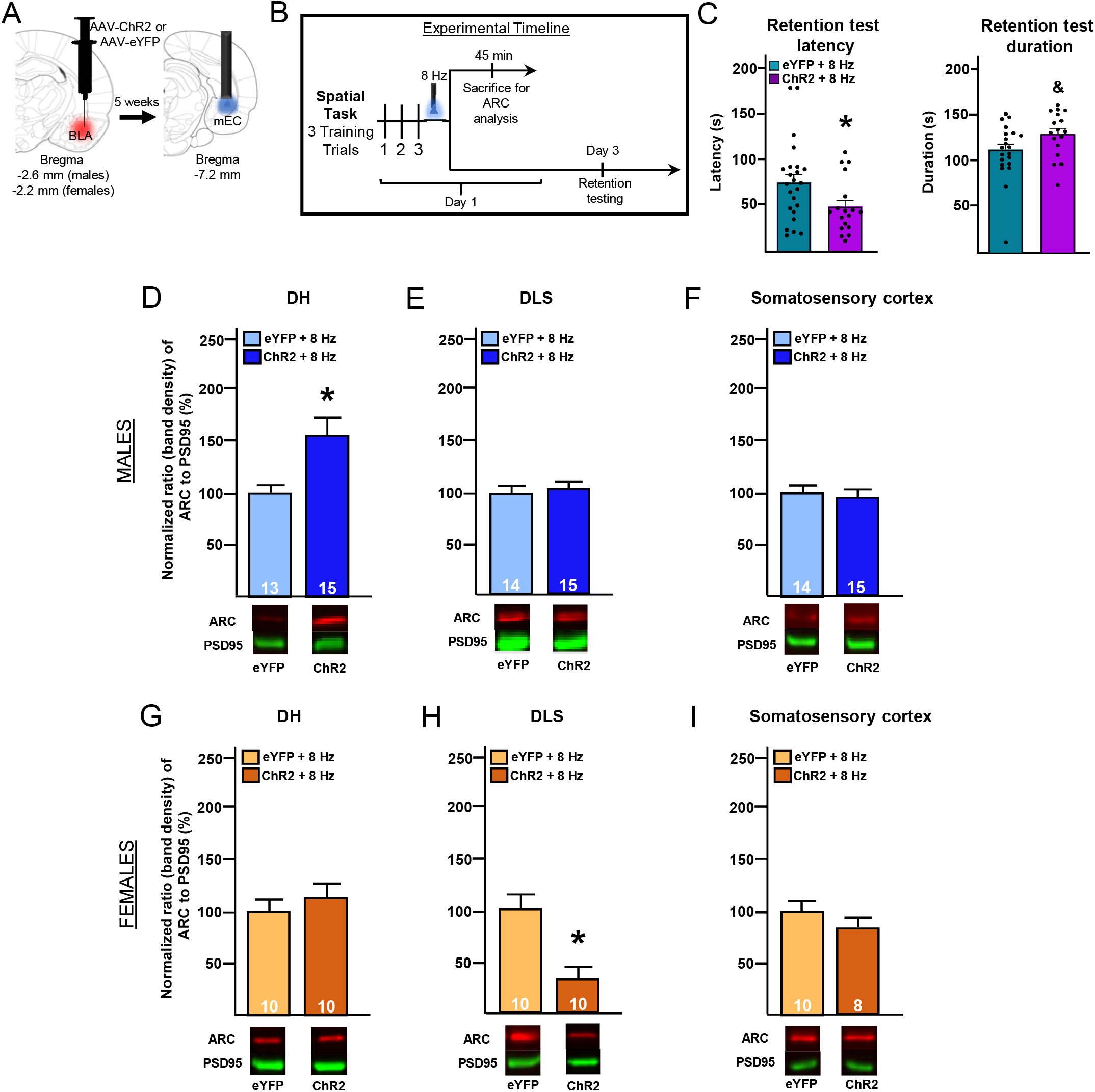
Effects of optical stimulation of the BLA-mEC pathway immediately after spatial training on ARC protein expression (Experiment 2). **A**, Schematic diagram of BLA injection site (left), incubation time, and optic probe placement in mEC (right). **B**, Experimental timeline for Experiment 2. **C**, Two days after training, half of the rats were tested for retention in a single 180 s trial. *Left*, Latencies to locate the escape port during the retention test. Rats that had received 8 Hz stimulation of ChR2-transduced BLA fibers in the mEC had significantly decreased latencies to find the escape port compared to their eYFP-control counterparts. *Right*, Duration spent in the target quadrant during the retention test. Rats that had received 8 Hz stimulation of the BLA-mEC pathway showed a trend towards increased time spent in the target quadrant of the maze compared to their eYFP-control counterparts. **D-F**, ARC protein expression (*top*) and representative immunoblots (*bottom*) in the DH (**D**), DLS (**E**), and somatosensory cortex (**F**) for male rats that were given spatial training immediately followed by 8 Hz stimulation of the BLA-mEC pathway. Rats that had received 8 Hz stimulation of the BLA-mEC pathway had significantly higher levels of ARC protein expression in the DH compared to eYFP-control animals. There were no significant differences in ARC expression in the DLS or somatosensory cortex. **G-I**, ARC protein expression (*top*) and representative immunoblots (*bottom*) in the DH (**G**), DLS (**H**), and somatosensory cortex (**I**) for female rats that were given spatial training immediately followed by 8 Hz stimulation of the BLA-mEC pathway. There were no significant differences in ARC expression in the DH or somatosensory cortex. However, rats that had received 8 Hz stimulation of the BLA-mEC pathway had significantly lower levels of ARC protein expression in the DLS compared to eYFP control animals. *, p < .05 compared to eYFP control values; &, p < 0.1, compared to eYFP control values. ARC is normalized to PSD95 by calculating the ratio band density of ARC to that of PSD95 and then expressed as a percentage of normalized control values.

#### Experiment 3 – Effect of BLA-mEC stimulation (8 Hz) alone on ARC expression

Experiment 3 examined whether stimulating the BLA-mEC pathway in the absence of behavioral training alters ARC protein expression, as previous studies indicate that ARC expression relies upon training (McIntyre et al., 2005). For this experiment, male and female rats received optical illumination of the mEC to provide illumination of either ChR2 or eYFP control-transduced BLA fibers in the absence of spatial training, using the same parameters as Experiment 2. Rats were sacrificed 45 min after stimulation (Figure 3A and 3B).

**Figure 3.**
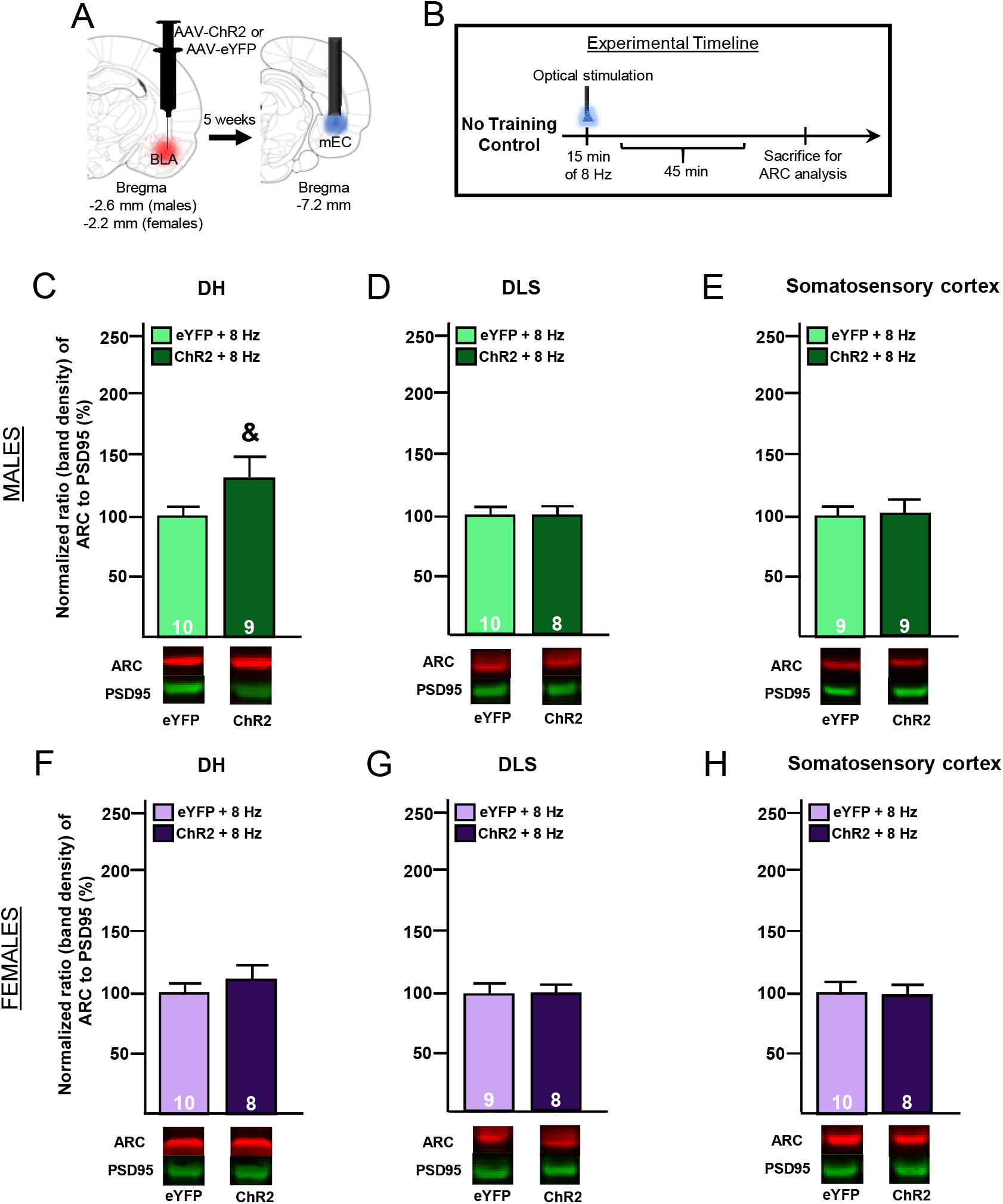
ARC protein expression effects of 8 Hz optical stimulation of the BLA-mEC pathway in the absence of behavioral training (Experiment 3). **A**, Schematic diagram of BLA injection site (left), incubation time, and optic probe placement in mEC (right). **B**, Experimental timeline for Experiment 3. **C-E**, ARC protein expression (*top*) and representative immunoblots (*bottom*) in the DH (**C**), DLS (**D**), and somatosensory cortex (**E**) for male rats that were given 8 Hz stimulation of the BLA-mEC pathway. Rats that had received 8 Hz stimulation of the BLA-mEC pathway had a trend toward significantly higher levels of ARC protein expression in the DH compared to eYFP control animals. There were no significant differences in ARC expression in the DLS or somatosensory cortex. **F-H**, ARC protein expression (*top*) and representative immunoblots (*bottom*) in the DH (**F**), DLS (**G**), and somatosensory cortex (**H**) for female rats that were given 8 Hz stimulation of the BLA-mEC pathway. There were no significant differences between groups in any case. &, p < 0.1, compared to eYFP control values. ARC is normalized to PSD95 by calculating the ratio band density of ARC to that of PSD95 and then expressed as a percentage of normalized control values.

#### Experiment 4 – Effect of BLA-mEC stimulation (40 Hz) alone on ARC expression

Based on the results with 8 Hz stimulation in male rats in Experiment 3, Experiment 4 examined whether stimulating the pathway with a different (higher) frequency would alter ARC expression. For this experiment, male rats received optical illumination of the mEC to provide illumination of either ChR2 or eYFP control-transduced BLA fibers in the absence of spatial training, using the following illumination parameters: 15 min of 2-s-trains of 40 Hz light pulses (pulse duration = 5 ms), given every 10 s (Wahlstrom et al., 2018). Rats were sacrificed 45 min after stimulation (Figure 4A and 4B).

**Figure 4.**
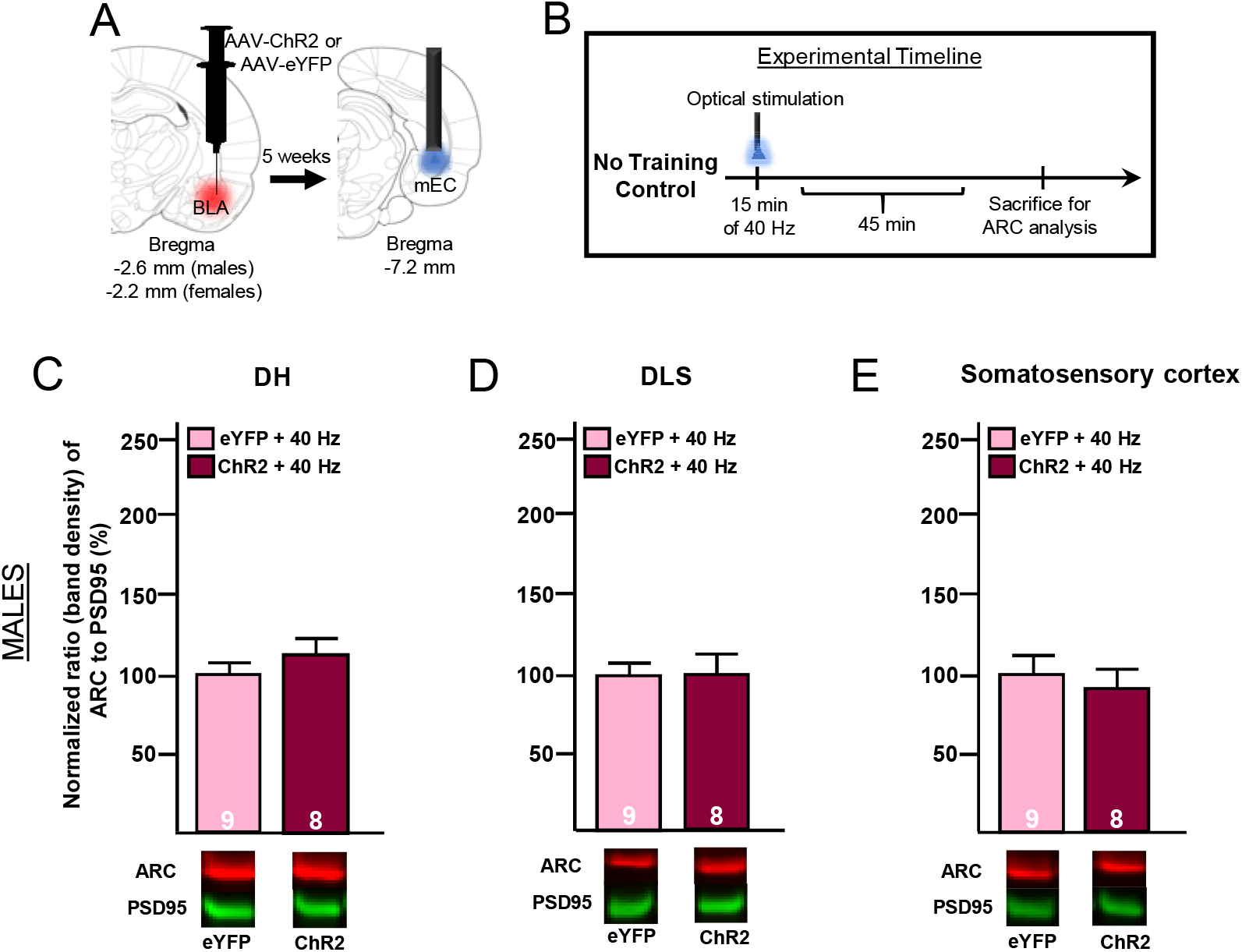
ARC protein expression effects of 40 Hz optical stimulation of the BLA-mEC pathway in the absence of behavioral training (Experiment 4). **A**, Schematic diagram of BLA injection site (left), incubation time, and optic probe placement in mEC (right). **B**, Experimental timeline for Experiment 4. Rats were given 15 min of 40 Hz optical stimulation of either ChR2 or eYFP control-transduced BLA axons in the mEC and sacrificed 45 min later for ARC protein analysis. **C-E**, ARC protein expression (*top*) and representative immunoblots (*bottom*) in the DH (**C**), DLS (**D**), and somatosensory cortex (**E**) for male rats that were given 40 Hz stimulation of the BLA-mEC pathway. There were no significant differences between groups in any case. ARC is normalized to PSD95 by calculating the ratio band density of ARC to that of PSD95 and then expressed as a percentage of normalized control values.

### Verification of opsin expression and histology

The following procedures were performed for rats used for ARC analysis that received virus microinjections and optic fiber implants. Rats were overdosed with sodium pentobarbital (200 mg/kg, i.p.), and brains were rapidly removed and flash-frozen in cold 2-methylbutane. The brains were coronally sectioned (75 μm) on a cryostat and mounted onto gelatin-subbed slides for ACEB (alkaline-stain-CAPS-ethanol-butanol) Nissl staining or mounted onto gelatin-subbed slides for immunohistochemical procedures. All slides were stored at −80° C until histological procedures began. Verification of optic probes’ placement was performed with an ACEB preparation (Lindroos and Leinonen, 1983) for rapid Nissl staining of frozen tissue and light microscopy according to the Paxinos and Watson atlas (Paxinos & Watson, 2014). Opsin expression in the BLA cell bodies was confirmed using immunohistochemistry procedures as described below. Slides were removed from the −80° C freezer and placed at 4° C for 20 min. Tissue sections were then fixed in ice-cold 4% paraformaldehyde for 60 min. Sections were soaked in 1x TBS and slides were removed one at a time for coverslipping with Vectashield HardSet Antifade Mounting Medium with DAPI (Vector Laboratories). eYFP expression was assessed by using a fluorescent microscope.

The following procedures were performed for rats used for retention testing that received virus microinjections and optic fiber implants. Animals were overdosed with sodium pentobarbital (100 mg/kg, i.p.) and transcardially perfused with phosphate-buffered saline (PBS) followed by PBS containing 4% paraformaldehyde. Brains were removed and stored at room temperature in 4% paraformaldehyde PBS for a minimum of 48 hours before sectioning. The brains were coronally sectioned (75 μm) on a vibratome and mounted onto either gelatin-subbed slides for Nissl staining or stored in anti-freeze solution at −20° C until immunohistochemical procedures began. Verification of optic probes’ placement was performed with a standard Nissl stain preparation (Cresyl violet) and light microscopy according to the Paxinos and Watson atlas (Paxinos & Watson, 2014). Expression in the BLA cell bodies and axons in the mEC were confirmed by using immunohistochemistry procedures as described below. Tissue sections were incubated in anti-GFP primary antibody solution for 48–72 h [PBS, 2% goat serum, 0.4% Triton X, rabbit 1:20,000 primary antibody (Abcam)]. Sections were then incubated for 1 h in a biotinylated anti-rabbit secondary antibody solution (K-PBS; 0.3% Triton X; goat, 1:200; Vector Labs) and incubated in an ABC kit (Vector Labs) for 1 h. Sections were developed in diaminobenzidine for ∼5–10 min before being mounted onto gelatin-subbed slides. Slides were allowed to dry before being dehydrated with reverse alcohol washes for 1 min each, soaked in Citrosolv for a minimum of 5 min, and coverslipped with DePeX (Electron Microscopy Sciences). eYFP expression was assessed by using either a light or fluorescent microscope.

### Statistical analysis

GraphPad Prism 8 was used for all statistical analyses in Experiments 1-4. For behavioral analysis, training latencies and time in target quadrant during training for Barnes maze experiments were analyzed using two-way ANOVAs. Retention latencies and durations in target quadrant for all behavioral experiments were analyzed using t tests. Scatter plots are included to best reflect the numerical spread of the data. For protein analysis, either a t test or a one-way ANOVA with a Holm-Sidak multiple comparisons test was used. The Grubbs method was used to identify statistical outliers across experiments. The α level was set to 0.05. All measures are expressed as mean ± SEM, and each group’s n is indicated in the figure below its respective bar.

## Results

### Experiment 1

Experiment 1 examined whether spatial or cued-response training in the Barnes maze task alters ARC protein expression. Figure 1A shows an illustration of the spatial version of the Barnes maze. Figure 1B shows an illustration of the cued-response version of the Barnes maze. Figure 1C shows the experimental timeline for Experiment 1. Figure 1D shows a schematic diagram illustrating the tissue punch sites where ARC was analyzed.

Two-way ANOVAs of spatial training latencies and duration in the target quadrant during training, respectively, revealed no significant difference between males and females (F_(1, 13)_ = 2.02, *p* = 0.18; F_(1, 13)_ = 1.05, *p* = 0.33; data not shown). Similarly, two-way ANOVAs of cued-response training latencies and duration in the target quadrant during training, respectively, revealed no significant difference between males and females (F_(1, 14)_ = 0.014, *p* = 0.91; F_(1, 14)_ = 2.67, *p* = 0.12; data not shown). Therefore, male and female rats were combined in each group.

Rats that underwent spatial training did not perform significantly differently on the 6^th^ and final training trial compared to rats that underwent cued-response training (mean spatial training latency ± SEM = 47.07 ± 4.25, mean cued-response training latency ± SEM = 47.69 ± 4.45); (mean spatial training duration ± SEM = 24.12 ± 4.42, mean cued-response training duration ± SEM = 24.49 ± 4.62). An unpaired t test of training latency and duration in target quadrant during training, respectively, revealed no significant difference between rats that received spatial training and rats that received cued-response training (t_(29)_ = 0.10, *p* = 0.92; t_(29)_ = 0.057, *p* = 0.95). Thus, rats that were trained on the spatial task did not show performance differences, compared to those rats that received cued-response training, on their respective tasks.

Figure 1E shows ARC protein expression in the DH for the three groups of rats. As males and females showed the same pattern of changes in ARC, males and females were not disaggregated. A one-way ANOVA revealed a statistically significant difference in ARC expression between groups (F_(2, 43)_ = 7.23, *p* = 0.002). Post hoc analyses indicated that ARC levels in rats that underwent spatial training was significantly higher compared to those observed in the no-training control group (*p* = 0.0015) and the cued-response group (*p* = 0.045). This is consistent with previous work showing increases in ARC expression in the DH of rats exposed to a spatial task (Guzowski et al., 2001; Ramirez-Amaya et al., 2005). However, ARC levels did not significantly differ between those rats given cued-response training and those in the no training-control group (*p* = 0.15).

Figures 1F and 1G show ARC protein expression in the DLS and somatosensory cortex respectively, for the three groups. One-way ANOVAs revealed no effect in either case (F_(2, 42)_ = 1.07, *p* = 0.35; F_(2, 42)_ = 1.14, *p* = 0.33;, respectively).

### Experiment 2

Experiment 2 examined whether stimulating the BLA-mEC pathway after spatial training alters ARC protein expression in the DH, DLS, and somatosensory cortex. Figure 2A shows a schematic diagram illustrating the site of virus injection into the BLA and the optical fiber implantation site aimed at the mEC. Figure 2B shows a timeline of behavioral training, optical stimulation, and ARC analysis or retention testing.

Two-way ANOVAs of spatial training latencies and duration in the target quadrant during training, respectively, revealed no significant difference between males and females that received post-training stimulation of ChR2 or eYFP control vector-transduced BLA axons in the mEC, respectively, (F_(1, 39)_ = 0.17, *p* = 0.69; F_(1, 39)_ = 2.21, *p* = 0.15; F_(1, 44)_ = 0.47, *p* = 0.50; F_(1, 44)_ = 0.38, *p* = 0.54). Thus, male and female rats did not differ from each other within each group and, as a result, were not disaggregated.

Rats that subsequently received post-training stimulation of ChR2-transduced BLA axons in the mEC on Day 1 did not show differences in their training on the final *training* trial compared to rats that subsequently received post-training stimulation of CaMKIIα-eYFP control vector-transduced BLA axons in the mEC on Day 1, (mean eYFP training latency ± SEM = 54.73 ± 1.83; mean ChR2 training latency ± SEM = 54.43 ± 1.90; mean eYFP training duration ± SEM = 16.14 ± 2.24; mean ChR2 training duration ± SEM = 19.78 ± 2.60). An unpaired t test of training latency and duration in target quadrant during training, respectively, revealed no significant difference between eYFP rats and ChR2 rats ((t_(85)_ = 0.12, *p* = 0.91); (t_(85)_ = 1.07, *p* = 0.29)). Thus, rats that received post-training stimulation of ChR2-transduced BLA axons in the mEC on Day 1 did not show differences in their training compared to control rats prior to optogenetic stimulation on Day 1.

Unpaired t tests of retention latencies and duration in target quadrant during *retention* testing, respectively, revealed no significant difference between males and females that received post-training illumination of ChR2 or CaMKIIα-eYFP control vector-transduced BLA axons in the mEC, (t_(16)_ = 0.45, *p* = 0.66; t_(16)_ = 0.15, *p* = 0.88; t_(20)_ = 0.55, *p* = 0.59; t_(20)_ = 0.48, *p* = 0.63). Therefore, male and female rats in each group were not disaggregated. Figure 2C shows retention latencies (left) and duration in target quadrant (right) for rats that were tested on Day 3. An unpaired t test revealed a significant difference in retention latencies (t_(40)_ = 2.22, *p* = 0.033) and a trend toward a significant difference in duration in target quadrant during retention testing (t_(38)_ = 1.94, *p* = 0.060). Consistent with our prior work, rats that had received 8 Hz post-training stimulation of BLA axons in the mEC required less time to find the escape port and spent more time in the target quadrant compared to eYFP-control rats.

The results from the analyses on ARC expression from male and female rats, shown in Figure 2D – Figure 2I, were disaggregated and the groups fully powered because the male and female rats appeared to differ in terms of effects of optical stimulation on ARC protein expression. Figure 2D shows ARC protein expression in the DH for male rats that were given spatial training immediately followed by 8 Hz stimulation of the BLA-mEC pathway. An unpaired t test revealed a significant difference between groups (t_(21.90)_ = 2.10, *p* = 0.048). Male rats that had received 8 Hz stimulation had significantly higher ARC protein levels in the DH compared to eYFP control rats. Figure 2E and 2F show ARC protein expression in the DLS and somatosensory cortex, respectively, for male rats given spatial training followed by 8 Hz stimulation of the BLA-mEC pathway. An unpaired t test revealed no significant difference between groups in either case (t_(27)_ = 0.33, *p* = 0.74; t_(27)_ = 0.43, *p* = 0.67, respectively).

Figure 2G and 2I show the ARC protein expression in the DH and somatosensory cortex, respectively, for female rats that were given spatial training immediately followed by 8 Hz stimulation of the BLA-mEC pathway. An unpaired t test revealed no significant difference between groups in either case (t_(18)_ = 0.89, *p* = 0.38; t_(16)_ = 0.68, *p* = 0.50, respectively). Figure 2H shows the ARC protein expression in the DLS for female rats that were given spatial training immediately followed by 8 Hz stimulation of the BLA-mEC pathway. An unpaired t test revealed a significant difference between groups (t_(18)_ = 2.38, *p* = 0.029). Rats that had received 8 Hz stimulation of ChR2-transduced BLA axons in the mEC had significantly lower levels of ARC protein expression in the DLS compared to eYFP-control rats.

### Experiment 3

Experiment 3 examined whether 8 Hz stimulation of the BLA-mEC pathway in the absence of spatial training alters ARC protein expression in the DH, DLS, and somatosensory cortex. Figure 3A shows a schematic diagram illustrating the site of virus injection into the BLA and the optical fiber implantation site aimed at the mEC. Figure 3B shows a timeline of optical stimulation and ARC analysis.

Figure 3C shows the ARC protein expression in the DH for male rats that were given 8 Hz stimulation of the BLA-mEC pathway in the absence of spatial training. Rats that had received 8 Hz stimulation of ChR2-transduced BLA axons in the mEC had higher levels of ARC protein expression in the DH compared to eYFP-control rats, though this only reached trend level (t_(17)_ = 1.99, *p* = 0.063). Figure 3D and 3E show the ARC protein expression in the DLS and somatosensory cortex, respectively, for male rats that were given 8 Hz stimulation of the BLA-mEC pathway in the absence of spatial training. An unpaired t test revealed no significant difference between groups in either case (t_(16)_ = 0.13, *p* = 0.90; t_(16)_ = 0.35, *p* = 0.73, respectively).

Figure 3F-3H show the ARC protein expression in the DH, DLS, and somatosensory cortex, respectively, for female rats that were given 8 Hz stimulation of the BLA-mEC pathway in the absence of spatial training. An unpaired t test revealed no significant difference between groups in any case (t_(16)_ = 1.24, *p* = 0.23; t_(15)_ = 0.031, *p* = 0.98; t_(16)_ = 0.011, *p* = 0.99, respectively).

### Experiment 4

Based on the results in Experiment 3 showing that 8 Hz optical stimulation alone alters ARC protein expression in the DH of male rats (but not female rats), Experiment 4 examined the frequency-specific nature of these effects, and whether 40 Hz stimulation of the BLA-mEC pathway in the absence of spatial training alters ARC protein expression in male rats. Figure 4A shows a schematic diagram illustrating the site of virus injection into the BLA and the optical fiber implantation site aimed at the mEC. Figure 4B shows a timeline of optical stimulation and ARC analysis.

Figures 4C-4E show the ARC protein expression in the DH, DLS, and somatosensory cortex, respectively, for male rats that were given 40 Hz stimulation of the BLA-mEC pathway in the absence of spatial training. An unpaired t test revealed no significant difference between groups in any case (t_(15)_ = 1.71, *p* = 0.11; t_(15)_ = 0.16, *p* = 0.88; t_(15)_ = 0.82, *p* = 0.43, respectively).

Table 1 summarizes the findings from the ARC analyses of all the experiments.

**Table 1.**
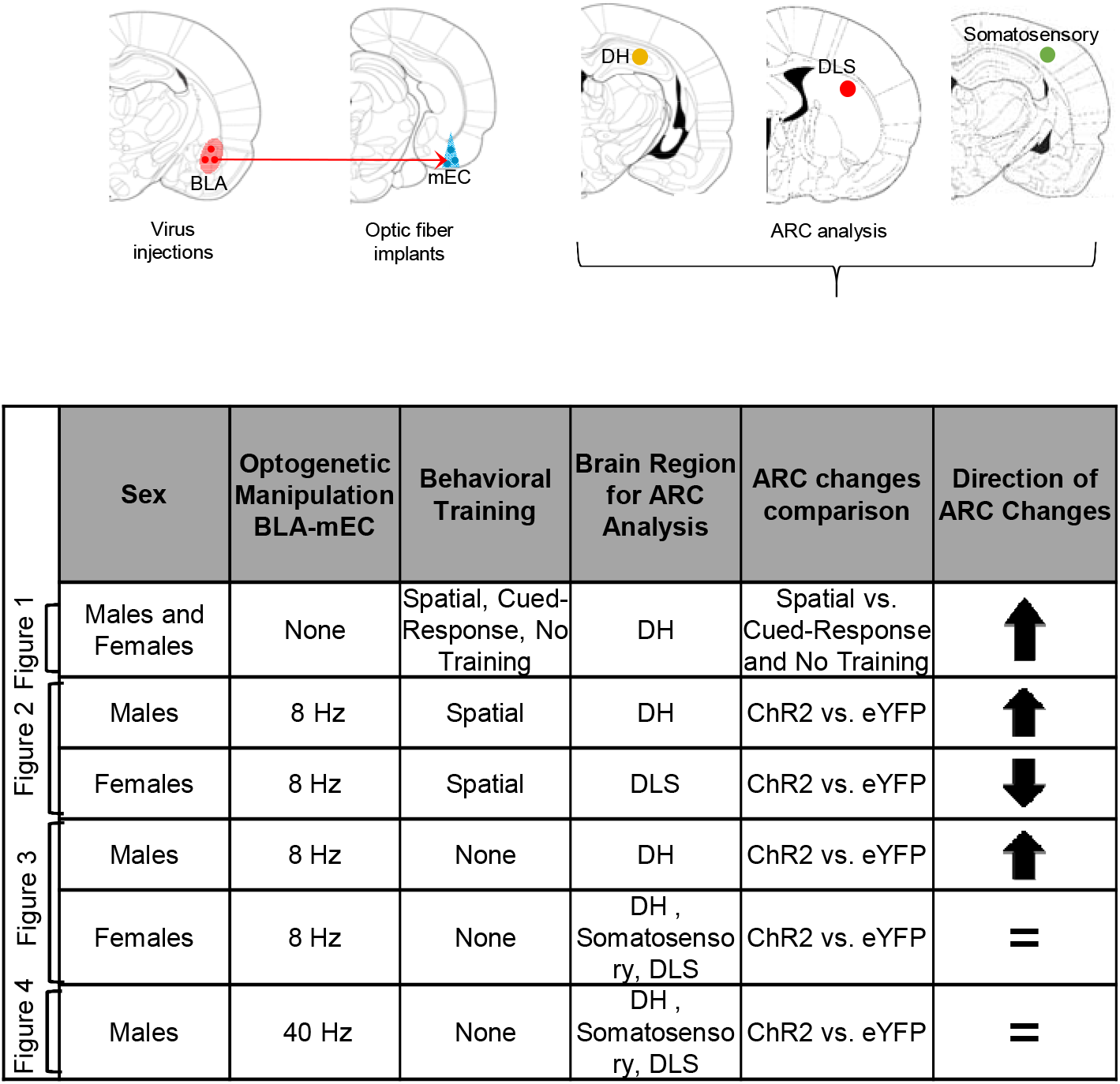
Summary of ARC findings from all Experiments. *Top*, Illustration of the BLA-mEC pathway that was optically manipulated in Experiment 2-4, followed by the brain regions analyzed for ARC protein (DH, DLS, and Somatosensory). *Bottom*, Summary of ARC expression findings across the four experiments conducted.

## Discussion

The current findings indicate a causal relationship between stimulating the BLA-mEC pathway and downstream changes in ARC expression, though this relationship appears to be more complex than expected. An initial experiment in the present study found that spatial training increased ARC levels in the DH of both male and female rats significantly above those found with cued-response training or no training. Subsequent experiments found that optogenetic stimulation of the BLA-mEC pathway immediately after spatial training significantly enhanced retention in males and females, confirming previous work using males alone from this laboratory (Wahlstrom et al., 2018). Such stimulation after training increased ARC protein levels in the DH with no effect on ARC expression in the DLS or somatosensory cortex, but only in male rats. In female rats, identical stimulation *reduced* ARC levels in the DLS without any effect on ARC levels in the DH or somatosensory cortex. This suggests that the ability of the BLA to modulate downstream ARC expression following spatial learning occurs, at least in part, through its projections to the mEC, though the precise region where ARC changes occur appears to depend on sex.

### The BLA-mEC pathway and downstream ARC expression

The present work found that spatial training in a Barnes maze increased ARC protein levels in the DH, but not DLS or somatosensory cortex, for male and female rats. This is consistent with prior work implicating ARC in hippocampal-dependent learning and memory (Guzowski et al., 2000; McIntyre et al., 2005; McReynolds et al., 2014). Indeed, evidence suggests that inhibition of ARC translation via antisense injections into the hippocampus impairs spatial memory and synaptic plasticity (Guzowski et al., 1999; Guzowski et al., 2000; McIntyre et al., 2005). Interestingly, the present study did not observe a similar increase in ARC with cued-response training in the Barnes maze, despite the overall similarity of the external environment. As cued-response and spatial learning involve the DLS and DH, respectively, this suggests that the combination of exposure to the external cues and the type of learning the rats engaged in was necessary for the observed increase in ARC in the DH. To our knowledge, this finding is also the first to show an effect on hippocampal ARC specific to the type of learning.

Much evidence suggests that ARC is a critical mechanism by which the BLA alters plasticity in other regions, as studies indicate that manipulations of BLA activity alter the expression of ARC protein downstream in the DH (McIntyre et al., 2005; McReynolds et al., 2014). However, it was unknown whether our prior work showing that stimulating the BLA-mEC pathway enhances retention for spatial training (Wahlstrom et al., 2018) also alters ARC expression in the DH. The present work first replicated the previous work from this laboratory, finding that stimulating this pathway with bursts of 8 Hz stimulation enhanced retention for spatial learning, in this case using both male and female rats. Immunoblotting analyses found that such stimulation following spatial training in male rats also increased ARC levels in the DH, but not DLS or somatosensory cortex.

However, such stimulation in male rats in the *absence* of spatial training also increased ARC in the DH. Most previous studies indicate that ARC expression is experience-dependent (Guzowski et al., 2001; Vazdarjanova et al., 2002; McIntyre et al., 2005; Ren et al., 2014). Indeed, posttraining intra-BLA infusions of the beta-adrenergic agonist clenbuterol enhance inhibitory avoidance retention and increase ARC levels in the DH but only do so when given after training (McIntyre et al., 2005). Nonetheless, the same study found that intra-BLA lidocaine infusions decreased ARC levels in the DH even in the absence of inhibitory avoidance training. Whether ARC expression can be altered in the absence of experience is unclear. One possibility in the current study is that simply placing the rats into a habituated context (the chamber where rats received optical manipulation), as was done in the stimulation-alone experiments, is sufficient to induce a small amount of ARC on its own and enable the stimulation to further increase ARC levels.

However, that 40 Hz stimulation of the BLA-mEC pathway without training in males had no effect on hippocampal ARC expression strongly suggests the answer may be more complex, as it appears that the stimulation parameters are a critical element in the ability of this pathway to alter hippocampal ARC. If non-specific BLA activation could alter hippocampal ARC expression, clenbuterol infusions as in the previous study (McIntyre et al., 2005) and 40 Hz stimulation alone in the present study should have altered ARC levels. The specificity of the 8 Hz stimulation is consistent with prior work indicating that such stimulation, but not 40 Hz stimulation, of the BLA-mEC pathway enhances the consolidation of spatial memories (Wahlstrom et al., 2018). Moreover, the prior work also found that stimulating the BLA-mEC pathway with 8 Hz pulses increases local field potentials at 8 Hz frequency bands in the DH, whereas similar stimulation with 40 Hz has much lower effects on local field potentials in the 40 Hz frequency. Much evidence suggests a critical role for theta rhythm (∼8 Hz) activity in the mEC and hippocampus in spatial information processing (Buzsaki, 2005; Buzsaki and Moser, 2013). Thus, the present findings add to the evidence that stimulating the BLA-mEC pathway in the same frequency range (i.e., ∼8 Hz) has privileged effects on DH activity, plasticity, and related memories.

### Sex differences in ARC expression

Although spatial training alone increased ARC levels in the DH of both sexes and BLA-mEC stimulation at 8 Hz enhanced retention for spatial learning in both sexes, such stimulation after spatial training did not alter ARC protein levels in the DH of female rats. Rather, such stimulation *decreased* ARC levels in the DLS of females. These results are surprising and yet, in light of the paucity of previous studies using both males and females, it is difficult to determine how unexpected they are. Although sparse, there have been reports of sex interactions with ARC expression (Henderson et al., 2017; Randesi et al., 2018; Randesi et al., 2019). Nonetheless, in each of those cases, the sex difference in ARC expression was one of degree, rather than kind. In contrast, the present experiments reveal a completely different and unexpected response in males vs. females.

That stimulation of the BLA-mEC pathway after training enhanced retention in both sexes while producing sex-dependent effects on ARC expression raises the possibility that these consequences on ARC levels represent different mechanisms for accomplishing the same outcome. Evidence suggests that males and females have biological differences yet often behave similarly (Gruene et al., 2015). This has led to the theory that some sex differences in the brain are compensatory and exist to equate behavior between males and females (De Vries, 2004; McCarthy et al., 2012). In this theory, sex convergence occurs when the endpoints are the same but the underlying physiology is different between sexes, an idea that may explain the present results.

One possibility consistent with this idea is that the changes in ARC in the DH and DLS in males and females, respectively, may reflect the competition that occurs between the hippocampus-based and basal ganglia-based memory systems (Packard and White, 1991; Packard et al., 1994; Poldrack and Packard, 2003; Wahlstrom et al., 2018). Studies indicate that stress and age differentially engage the hippocampus vs. the striatum in males and females (Kim et al., 2001; Schwabe et al., 2007; Konishi et al., 2013). Prior work also suggests that, when given the choice between place and response strategies, female rats in estrous are biased towards response strategies (Korol et al., 2004). Thus, in the present experiments, optogenetic stimulation may have enhanced spatial memory in male rats by increasing ARC levels in the DH and in female rats by decreasing ARC levels in the competing memory system, the DLS.

Considering that spatial training alone increases ARC in the DH in both sexes, this raises the question of why BLA-mEC stimulation does not further increase ARC in the DH of females. It is possible that, in females, a “ceiling” effect prevented the stimulation from further increasing ARC in the DH above that obtained with spatial training. As stimulation alone in female rats did not increase ARC, however, a ceiling effect seems unlikely. Rather, that BLA-mEC stimulation after training decreased ARC in the DLS in females suggests that the stimulation alters the balance between the two memory systems. Alternatively, the changes in ARC expression in both males and females may be an ancillary consequence of stimulation that is unrelated to memory consolidation. This is unlikely at least for the effects on ARC in the DH in males, as considerable evidence points to a functional role of such ARC in consolidation (Guzowski et al., 2000; Holloway and McIntyre, 2011; McReynolds et al., 2014). However, whether changes in ARC expression in the DLS influences memory consolidation for either males or females is unknown. Nonetheless, these findings support the overall notion that the BLA-mEC pathway is an important mechanism by which the BLA influences synaptic plasticity in downstream brain regions.

## Acknowledgements

The authors declare no competing financial interests. This work was supported by NIH grants MH118754 (KLW) and MH104384 (RTL and CKM).

The authors thank John Wemmie, Rong Fan, and Joshua Weiner for their excellent technical assistance and for allowing us to utilize their laboratory facilities for the experiments involving Western blotting.

